# Galectin-1 Modulates Hepatocellular Carcinoma Response to Thermal Ablation Through Regulating Glycolysis

**DOI:** 10.1101/2024.12.12.628238

**Authors:** Tu Nguyen, Yonghwan Shin, Janet Pham, Aravinth Ruppa, Po-Chun Chen, Hannah Mirmohammadi, David S. K. Lu, Steven S. Raman, Jason Chiang

**Author notes:** **Conflict of interest**: All authors declare no conflict of interest. **Financial support statement**: This work was supported by the following grants: UCLA Departmental Exploratory Research Grant; UCLA Jonsson Comprehensive Care Center Fellowship Award. **Author contributions**: Conceptualization, T.N., J.C.; Methodology, T.N., J.C., Investigation, T.N., J.P., Y.S., A.R., and P.C.; Writing—Original Draft, T.N., J.C.; Writing—Review & Editing, T.N., J.C., Y.S., D.L., and S.R.; Data Analysis, T.N, J.C.; Supervision, T.N., J.C.; Funding Acquisition, T.N., J.C.; Resources, D.L., S.R., and J.C.

## Abstract

I.

**Background & Aims:** Thermal ablation is the standard of care treatment modality with curative intent for early-stage hepatocellular carcinoma (HCC), but its efficacy remains moderately limited— with up to 40% of HCC patients experiencing local recurrence post-treatment. This study aimed to evaluate the prognostic value of galectin-1 (Gal-1) in predicting thermal-ablation responsiveness. We then evaluated the therapeutic potential of targeting Gal-1 in inhibition of glycolysis and subsequently enhancing thermal-ablation efficacy.

**Methods:** Liquid-Chromatography Mass-Spectrometry (LC-MS) was employed to analyze proteomic profiles of retrospectively collected pre-ablation FFPE samples of known thermal-ablation responders and nonresponders. An *in-vitro* thermal peri-ablation model was established using a heated water bath. Gal-1 inhibition via OTX008 or knockdown was utilized to investigate hyperthermic sensitivity. Hyperthermia-resistant SNU449 cells were used to establish an orthotopic murine model to evaluate the combination therapy of OTX008 and thermal ablation. Harvested tumors were analyzed by LC-MS to determine their metabolic profiles.

**Results:** This study revealed that responders had significantly longer tumor progression-free survival compared to nonresponders (57.0±1.6 (median not reached) versus 8.3±0.5 months (median: 13.6 months), p<0.001). Moreover, responders were found to have significant downregulation of Gal-1 expression compared to that of nonresponders. Gal-1 inhibition or knockdown markedly increased hyperthermic sensitivity in hyperthermia-resistant HCC SNU449 cells. Targeting Gal-1 by OTX008 in combination with thermal ablation significantly reduced SNU449-derived tumor growth compared to the thermal-ablation alone group *in vivo*. Metabolomic analysis revealed decreased glycolytic metabolites, fructose 1,6-bisphosphate, 3-phosphoglycerate and phosphoenolpyruvate, while western blot analysis showed decreased Gal-1 expression in the combined treatment group compared to monotherapy thermal ablation or OTX008 treatment.

**Conclusions:** Gal-1 overexpression correlates with thermal-ablation nonresponsiveness, and targeting Gal-1 enhances thermal-ablation efficacy by inhibiting glycolysis.

**Impact and Implications:** Despite being a standard-of-care treatment for early-stage HCC, thermal ablation has a high local recurrence rate of approximately 40%. While thermal ablation can lead to cellular death in the central-treatment zone, its metabolic impact on cells in the peri-ablational region remains unclear. This study shows the direct association between Gal-1 overexpression and thermal-ablation nonresponsiveness. Moreover, it found that Gal-1 inhibition or knockdown increased hyperthermia sensitivity *in vitro*. Targeting Gal-1 in combination with thermal-ablation significantly reduced hyperthermia-resistant SNU449 tumor growth by inhibiting glycolysis *in vivo*. These findings suggest that the efficacy of thermal ablation in HCC can be enhanced by pharmacologically inhibiting Gal-1.

**Graphical abstract:** 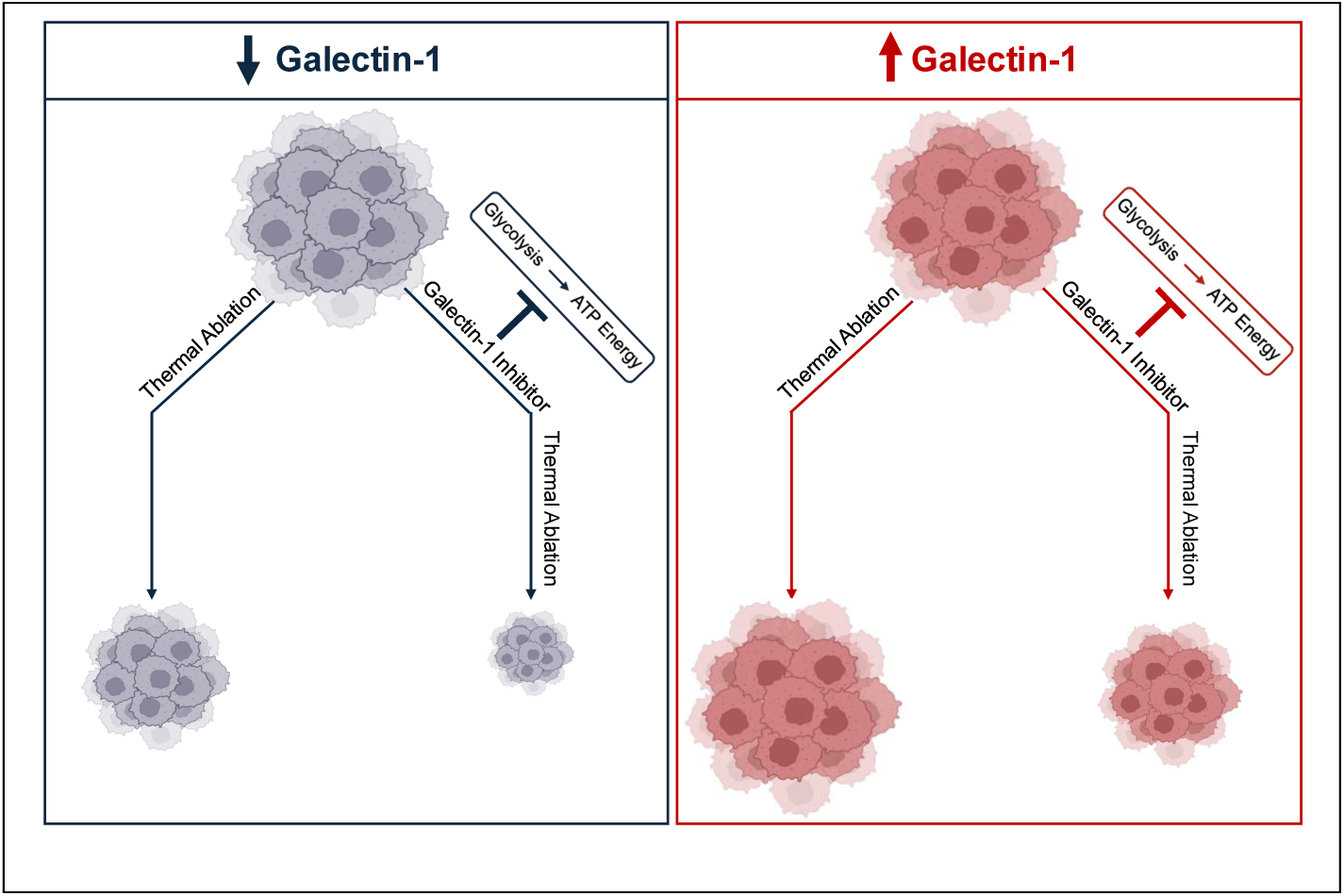

**Highlights:**

- Profiling pre-ablation HCC biopsies reveals Galectin-1 as a key prognostic biomarker for response prediction in thermal ablation
- Targeting Galectin-1 with a selective inhibitor (OTX008) enhances the efficacy of thermal ablation in HCC
- Galectin-1 modulates thermal-ablation response via regulating glycolysis in HCC

## II. INTRODUCTION

Hepatocellular carcinoma (HCC) is the most common primary liver cancer and the third leading cause of cancer-related death in the world^1^. Thermal ablation has historically been utilized with curative intent for early-stage and non-surgical HCC patients^2^. While the goal of thermal ablation is to treat the targeted HCC with adequate margins to achieve complete pathologic necrosis, there are select cases where there is rapid HCC progression after ablation, even in cases with technical success with adequate margins^3,4^.

The rationale for rapid HCC progression lies in the fact that sublethal hyperthermia is created at the edges of the ablation zone^5^. This sublethal hyperthermic environment is insufficient to eliminate tumor cells and can create a pro-tumorigenic milieu, exacerbating metabolic derangements associated with more aggressive HCC phenotypes^5^. These metabolic interactions can be amplified in HCC tumors that are more likely to have imaging-occult disease, such as in poorly-differentiated or macro trabecular subtypes, when exposed in the hyperthermic regions of the ablation zone^6^.

Metabolic alterations within the tumor microenvironment is known to play a key role in HCC treatment resistance and progression^7^, as rapid growth requires increasing energy production and macromolecular biosynthesis. However, in-depth analysis of early-stage HCC tumor microenvironment and associated metabolic derangements, particularly under hyperthermic conditions, has not been well evaluated. Part of the reason for this knowledge gap is that studying the molecular aberrations of HCC require clinical tissue samples for analysis—which is currently not within standard guidelines for HCC diagnosis^2,8^. HCC management guidelines recommend diagnosis of HCC based on imaging alone, restricting the use of needle biopsy in only select equivocal cases rather than routine diagnosis^2^. As a result, the molecular mechanism behind early-stage HCC progression and association with metabolic signal transduction pathways remains understudied.

A potential target that has been implicated in HCC progression is galectin-1 (Gal-1), an evolutionarily conserved, glycan-binding protein that has been associated with cancer aggressiveness and treatment resistance^9,10^. Gal-1 has been shown to be overexpressed in other different tumor types such as prostate and glioblastomas^11,12^. Additionally, Gal-1 has been found to regulate aerobic glycolysis—a key ATP-producing mechanism seen across many solid cancer types, also known as the Warburg effect^13,12^. However, Gal-1 has not yet been linked to regulation of HCC metabolism and promoting post-treatment resistance. This study aims to elucidate the role of Gal-1 in modulating HCC response to thermal ablation through glycolysis. In particular, we used liquid chromatography-mass spectrometry to identify Gal-1 differential expression in a unique retrospective set of pre-ablation needle-biopsy samples from early-stage HCCs, who were later on identified as thermal-ablation responders and nonresponders. Gal-1 modulation was shown to directly correlate with hyperthermia sensitivity *in vitro*. Finally, targeting Gal-1 was found to enhance the efficacy of thermal ablation by inhibiting glycolysis in a hyperthermia-resistant HCC mouse orthotopic model.

## III. RESULTS

### GALECTIN-1 OVEREXPRESSION CORRELATES WITH THERMAL-ABLATION NONRESPONSE IN HCC FFPE NEEDLE BIOPSY SAMPLES AND *IN VITRO*

Most HCC patients experience an initial complete response after ablation, but up to 40% of patients will eventually progress **(Fig.1A)**^3,4^. This is because sublethal hyperthermia associated with the peripheries of the thermal ablation zone **(Fig.1B)** is insufficient to eradicate surrounding imaging-occult tumor cells^5^. The Kaplan-Meier analysis of patients with matched-propensity scores **(Table 1)** showed improved outcomes of patients who were considered thermal-ablation responders (n=34), defined as those who had a complete response by localized mRECIST criteria for up to two years, compared to thermal-ablation nonresponders (n=24), defined as patients who had locally progressive disease by localized mRECIST criteria within the two years. Responders were shown to have significantly longer tumor progression-free (PFS) survival compared to nonresponders (57.0±1.6 (median not reached) versus 8.3±0.5 months (median: 13.6 months),p <0.001) **(Fig.1C)**. Additionally, the one and three-year PFS rates were significantly higher in responders versus nonresponders: 97.1% versus 41.0% and 81.1% versus 0.0% (p<0.001), respectively.

**Figure 1:**
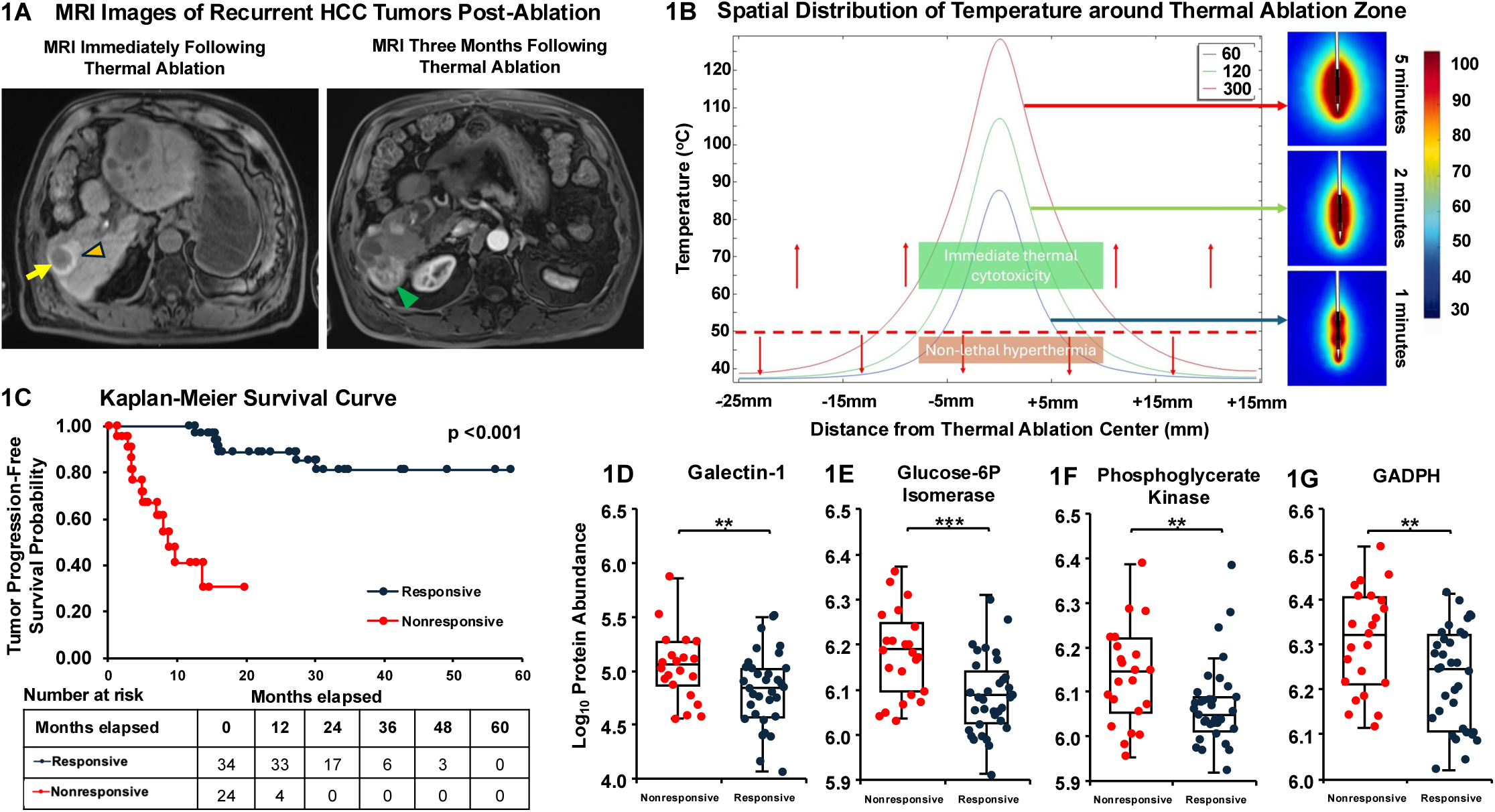
Upregulation of galectin-1 and critical glycolytic enzymes correlates with nonresponsivess to thermal ablation in HCC. **A)** (Left) T1-weighted non-contrast MRI of an HCC patient showing a completely ablated HCC tumor (yellow arrow) and adequate margins (orange arrowhead) around the target. (Right) Three-month follow-up post-contrast-arterial phase MRI demonstrating new growth (green arrowhead) around the prior-ablation zone. **B)** Ablation temperature distribution illustrating spatial temperature distribution from the radial center of a thermal-ablation zone. Note the peripheral one corresponds to the regions exposed to sublethal hyperthermia. **C)** Kaplan-Meier analysis of tumor progression-free survival probability in responders versus nonresponders. **D-G)** Upregulated levels of Galectin-1 **(D)**, Glucose-6-Phosphase Isomerase **(E)**, Phosphoglycerate Kinase **(F)**, Glyceraldehyde 3-Phosphate Dehydrogenase (GADPH) **(G)**, HCC responders (n=34) and nonresponders (n=24). Intensities were log10-transformed. p-value was calculated using one-tailed-unpaired Student’s T-test with **p<0.01, ***p<0.001. Box plots show median (central line), upper and lower quartiles (box limits), and 1.5 interquartile range (whiskers).

**Table 1:** Demographic and clinical characteristics of study patients. Continuous variables with normal distribution, including age, BMI (body mass index), tumor size, were shown as mean ± standard deviation. P-values were calculated using unpaired student’s t-test and chi-square for continuous variables and categorical variables, respectively. HCC, (hepatocellular carcinoma); HBV, hepatitis B; HCV, hepatitis C; NASH, nonalcoholic steatohepatitis. * indicates Child-Pugh B and C. ** indicates other immunocompromised states for the nonresponder cohort.

| Characteristics | Responders<br>(n=34) | Nonresponders<br>(n=24) | p-value |
| --- | --- | --- | --- |
| Age (year) | 68.8 ± 10.0 | 64.9 ± 10.2 | 0.159 |
| Male sex (%) | 23 (67.7%) | 18 (75.0%) | 0.545 |
| BMI (kg/m<sup>3</sup>) | 24.4 ± 5.5 | 24.8 ± 4.9 | 0.784 |
| Ablation modality | 34 (100% MWA) | 24 (100% MWA) |  |
| Child-Pugh Class % |  |  | 0.376 |
| A | 32(94.1%) | 21 (87.5%) |  |
| Others* | 2 (5.9%) | 3 (12.5%) |  |
| Etiology of HCC |  |  |  |
| HBV | 8 (23.5%) | 6 (25.0%) | 0.897 |
| HCV | 13 (38.2%) | 6 (25.0%) | 0.290 |
| NASH | 3 (8.8%) | 5 (20.8%) | 0.034 |
| Alcohol | 3 (8.8%) | 4 (16.7%) | 6.372x10<sup>-4</sup> |
| Others** | 7 (20.6%) | 3 (12.5%) | 0.156 |
| Multifocal % | 14 (41.2%) | 13 (62.5.0%) | 0.110 |
| Tumor size (cm) | 2.9 ± 1.1 | 2.8 ± 1.6 | 0.692 |
| Tumor lobe % |  |  |  |
| Left | 12 (35.3%) | 6 (25.0%) | 0.404 |
| Right | 21 (61.8%) | 14 (58.3%) | 0.792 |
| Both | 1 (2.9%) | 4 (16.7%) | 0.067 |
| Histopathological grade % |  |  |  |
| Well-differentiated | 1 (2.9%) | 0 (0%) | 0.397 |
| Moderately-differentiated | 31 (91.2%) | 21 (87.5%) | 0.651 |
| Poorly-differentiated | 1 (2.9%) | 3 (12.5%) | 0.157 |
| **Table 1:** Demographic and clinical characteristics of study patients. Continuous variables with normal distribution, including age, BMI (body mass index), tumor size, were shown as mean ± standard deviation. P-values were calculated using unpaired student's t-test and chi-square for continuous variables and categorical variables, respectively. HCC, (hepatocellular carcinoma); HBV, hepatitis B; HCV, hepatitis C; NASH, nonalcoholic steatohepatitis. * indicates Child-Pugh B and C. ** indicates other immunocompromised states for the nonresponder cohort. | | | |

Proteomic analysis was performed on respectively-collected FFPE biopsy samples, which showed that there was a significant elevation of Gal-1 in nonresponders compared to responders (p=0.0024) **(Fig.1D)**. Furthermore, this analysis revealed critical downstream proteins involved in the aerobic glycolysis—a key mechanism of ATP production conserved across many other solid cancer types, also known as the Warburg effect^13^—were also markedly upregulated in nonresponders: glucose-6-phosphate isomerase (p=0.00017), phosphoglycerate kinase (p=0.0036), and glyceraldehyde-3-phosphate dehydrogenase (GADPH) (p=0.0048) **(Fig.1E-G)**.

An *in-vitro* thermal peri-ablation model was created to simulate the responder-and-nonresponder model to further investigate the role of Gal-1. HCC-derived cell lines (SNU423, SNU449, and HepG2/C3a)^14,15^ were exposed to either normothermic (37°C) or sublethal hyperthermic (47°C) temperatures in a water bath for 10 minutes after an equilibrating period of 15 minutes **(Fig.S1A-B)**. Consistent with prior studies simulating the peri-ablational zone^16–18^, sublethal temperature defined as 47°C, was used to induce partial cell death without necrosis, preserving the ability to investigate underlying metabolic changes. The cell death percentage under hyperthermia was then determined in relation to respective controls at 24, 48, and 72 hours. SNU449 cells were found to be significantly more thermally resistant, with a markedly lower cell-death percentage under hyperthermic 47°C than SNU423 throughout the study periods **(Fig.2A-B)**. Moreover, this cell growth reduction pattern was confirmed in bright-field microscopy images taken at 24, 48, and 72 hours **(Fig.S1C)**, confirming SNU423 and SNU449 as hyperthermia-responsive and resistant cells, respectively. The relation of Gal-1 to sublethal hyperthermia, previously noted in the proteomic analysis of clinical FFPE samples, was then evaluated in this *in-vitro* model. Western blots confirmed Gal-1 upregulation in hyperthermia-resistant SNU449 cells compared to hyperthermia-responsive SNU423 cells **(Fig.2C)**. These findings suggest a link between Gal-1 levels and the response to thermal stress in both HCC patients and cell lines.

**Figure 2:**
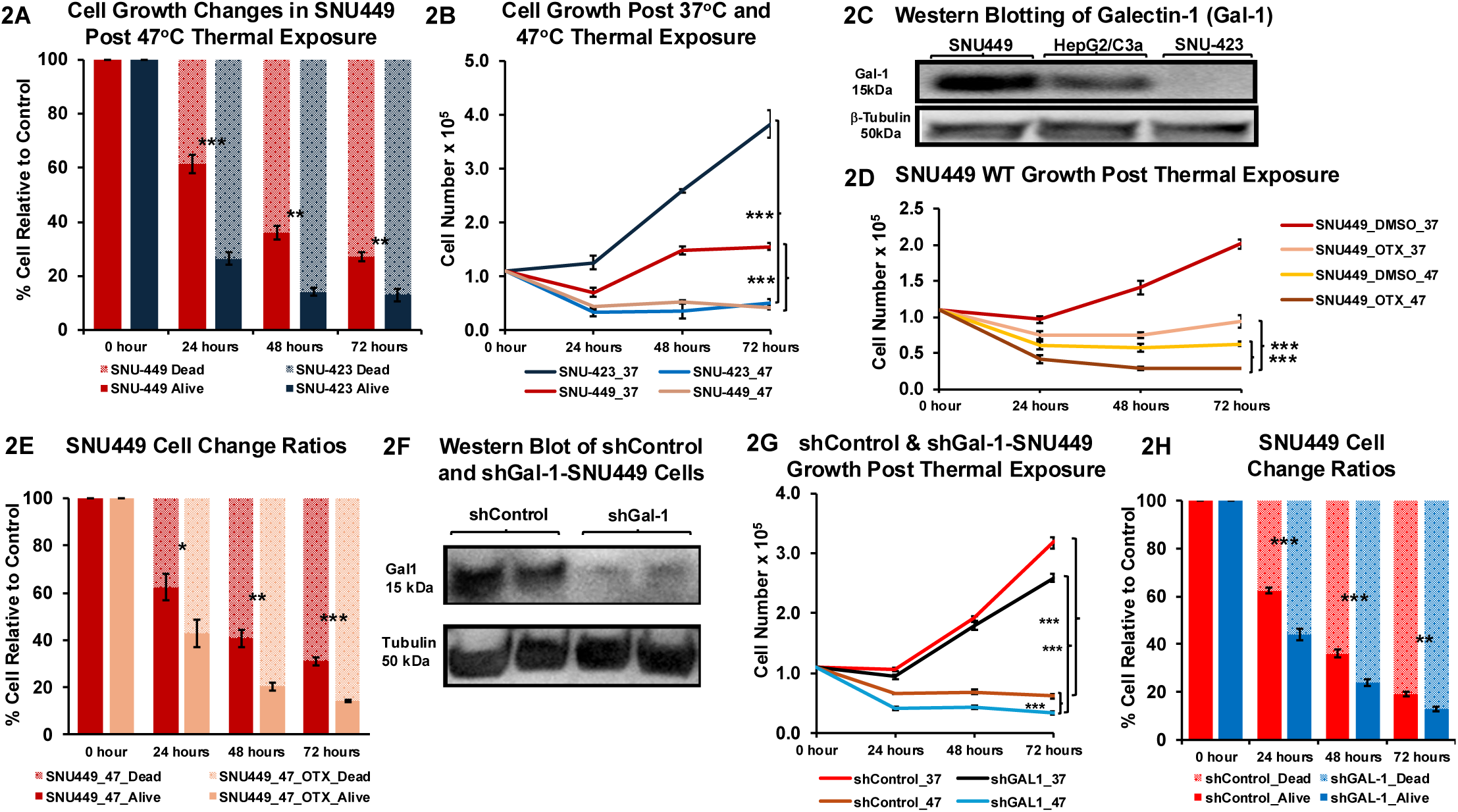
Galectin-1 overexpression is associated with sublethal-hyperthermia resistance. **A-B)** SNU449 survival rates (n=3 each) **(A)** post-thermal exposure at 37°C and 47°C at 24, 48, and 72 hours. Corresponding cell growth **(B)**. **C)** Western blot of Gal-1 in SNU449, SNU423, and HepG2/C3a (positive control). β-Tubulin was used to as loading control. **D-E)** SNU449 cell growth (n=3 each) **(D)** post-thermal exposure at 37°C and 47°C with Gal-1 inhibitor OTX or DMSO at 24, 48, and 72 hours. Corresponding cell survival rates **(E)**. **F)** Western blot of Gal-1 in shGal-1-SNU449 (SNU449 with Gal-1 knockdown) and respective shControl-SNU449. **G-H)** shControl and shGal-1-SNU449 cell growth (n=3) **(G)** post-thermal exposure at 37°C and 47°C at 24, 48, and 72 hours. Corresponding cell survival rates **(H)**. p-values were calculated using one-tailed-unpaired student’s t-test, *p<0.05, **p<0.01, ***p<0.001.

## GALECTIN-1 OVEREXPRESSION IS ASSOCIATED WITH SUBLETHAL HYPERTHERMIA RESISTANCE IN HCC CELLS

Given the finding of Gal-1 overexpression in hyperthermia-resistant SNU449 **(Fig.2C)**, selective modulation of Gal-1 expression was performed to isolate the contributory role of Gal-1 in promoting resistance to sublethal hyperthermia seen in the peri-ablation zone. SNU449 was treated with pre-determined 50µM Gal-1 inhibitor OTX^19^ **(Fig.S2A-B)** prior to being exposed to 37°C or 47°C. SNU449 cell growth with OTX treatment under hyperthermia was markedly diminished compared to that of those only exposed to 47°C **(Fig.2D)**. Also, cell death percentage in SNU449 demonstrated that silencing Gal-1 expression significantly increased their sensitivity to hyperthermia **(Fig.2E)**. An increase in hyperthermia response was similarly noted when Gal-1 was inhibited using OTX in SNU423 **(Fig.S2C-D)**.

Gal-1 silencing using lentiviral particles carrying shRNA was also used to reduce Gal-1 expression in SNU449 **(Fig.2F)**. The effect of silencing Gal-1 in shGal-1-SNU449 along with respective shControl-SNU449 was evaluated under hyperthermia. The cell growth indicated that shGal-1-SNU449 was significantly more susceptible to sublethal hyperthermia than shControl-SNU449 **(Fig.2G)**. The cell survival percentage analysis corroborated the cell growth results **(Fig.2H)**. These findings highlight that reducing Gal-1 expression enhances HCC cells’ sensitivity to hyperthermia.

## TARGETING GALECTIN-1 BY OTX008 ENHANCES EFFICACY OF THERMAL ABLATION IN HCC TUMORS THROUGH INHIBITING GLYCOLYSIS

Sublethal hyperthermia-resistant SNU449 cells were used to evaluate the efficacy of Gal-1 inhibition in improving post-ablation tumor control *in vivo* orthotopic murine model. This cell line was used for the *in-vivo* studies because success in controlling tumor progression with this hyperthermia-resistant cell line would further emphasize the critical role of Gal-1 in enabling these cells to develop metabolic plasticity for survival. After subcutaneous SNU449-cell-derived tumors were orthotopically implanted into the murine livers, each subject was administered with selective Gal-1 inhibitor OTX prior to thermal-ablation treatment with a timeline as shown **(Fig.3A)**. In accordance with ethical guidelines on animal well-being and appearance, recommended by the Division of Laboratory Animal Medicine at UCLA, mice were sacrificed one-week post-ablation, and tumors were harvested. Tumor weights were found to be significantly lower in the group that received both thermal ablation and the selective Gal-1 inhibitor OTX treatments, compared to thermal ablation or OTX alone **(Fig.3B-C)**. There was no reduction in tumor growth in the group that received thermal ablation alone compared to the control group, which was expected because SNU449 cells have been shown previously to be resistant to thermal ablation **(Fig.2A-B**, S1C). Also, there was no reduction in tumor growth observed in the selective Gal-1 inhibitor OTX alone group compared to the control **(Fig.3B-C)**.

**Figure 3:**
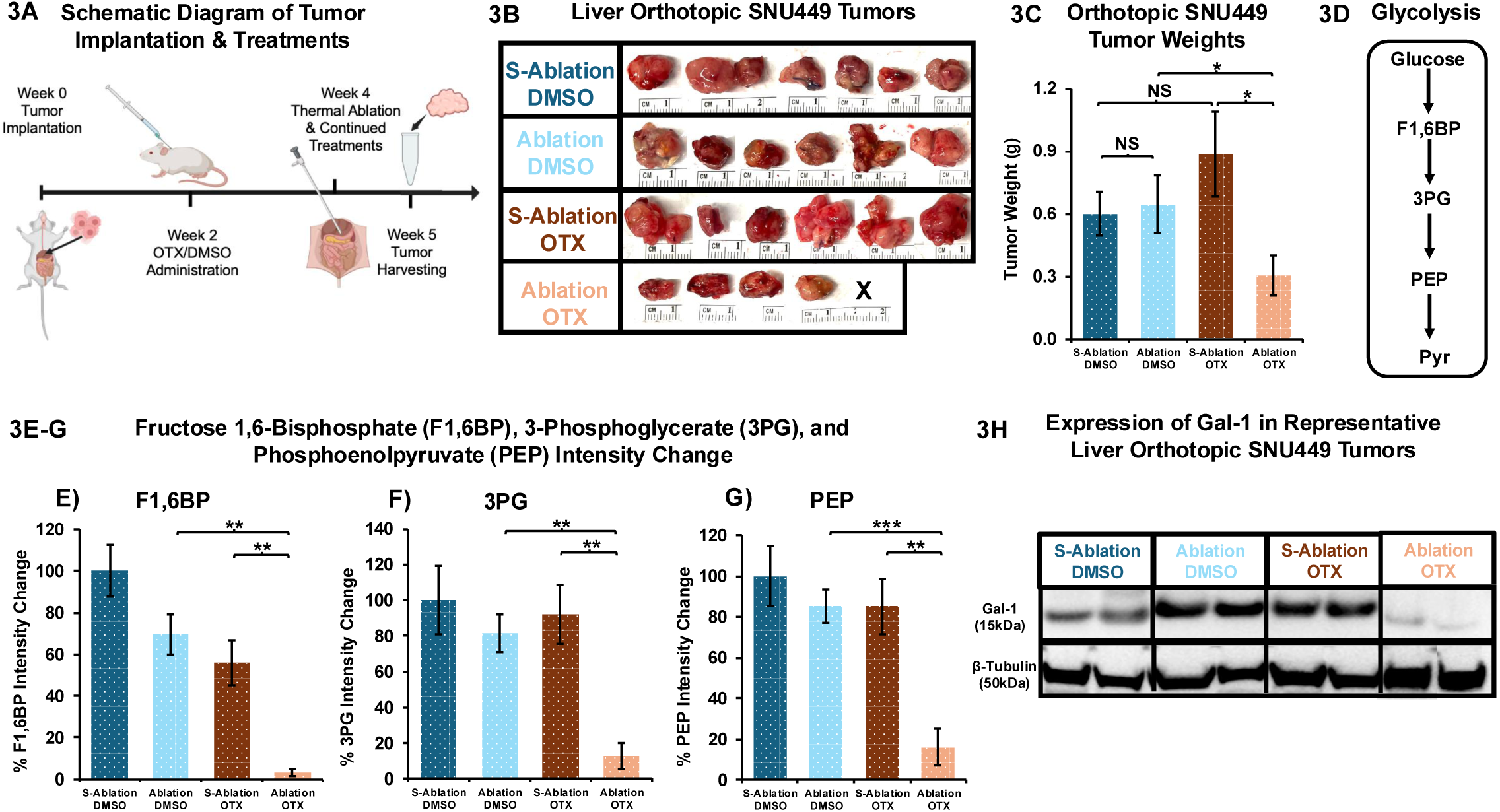
Targeting galectin-1 enhances efficacy of thermal ablation through inhibiting glycolysis. **A)** Schematic diagram showing experimental timeline of events. **B)** Tumors from the study groups: S-ablation DMSO (S=sham) (n=6), ablation DMSO (n=6), S-ablation OTX (n=6), ablation OTX (n=5, X indicates tumor disappearance). **C)** Respective tumor weights. **D)** Schematic diagram of glycolysis. **E-G)** Intensity change for glycolytic metabolites in respective tumors: fructose-1,6-bisphosphate **(E)** 3-phosphoglycerate **(F),** phosphoenolpyruvate **(G)**. Percent change was compared to respective controls. **H)** Western blot Gal-1 expression in respective tumors. β-Tubulin was used as loading control. p-values were calculated using one-tailed-unpaired student’s t-test, *p<0.05, **p<0.01, ***p<0.001.

The altered metabolic profile of each group was then characterized. In particular, the analysis focused on glycolysis due to the clinical findings of upregulations of critical glycolytic proteins found in FFPE biopsy specimens **(Fig.1E-G)**. When mass spectrometry was employed to profile glycolytic metabolites **(Fig.3D)**, fructose 1,6-bisphosphate (F1,6BP), 3-phosphoglycerate (3PG), and phosphoenolpyruvate (PEP) levels were found to be consistently diminished in the combined treatment group compared to thermal ablation or OTX alone **(Fig.3E-G).** These findings confirm a significant reduction in glycolysis in the combined group and corroborated the clinical findings of downregulation of glycolytic activities in thermal-ablation responsive group **(Fig.1E-G)**. Additionally, Gal-1 levels in the harvested tumors were assessed, where it was noted that Gal-1 levels in the combined treatment group was significantly reduced as compared to any other study group **(Fig.3H)**. This finding is in concordance with current literature describing degradation of OTX bound-Gal-1 complex being degraded by hyperthermia-exposed 26S proteasome^20,21^. Collectively, the findings highlight the critical role of Gal-1 in modulating thermal-ablation response via regulating glycolysis.

## IV. MATERIALS/PATIENTS AND METHODS

### Patient cohorts

This retrospective study was performed under written-informed consents and approval from UCLA Institutional Board Review (IRB#23-000131). A total of n=58 previously-banked biopsies were obtained from patients who had been diagnosed with moderate-differentiated (n=54) or poorly-differentiated (n=4) HCC. Thermal-ablation responsiveness was monitored for up to 5 years (60 months). Responders (n=34) were defined as patients who had complete response by localized mRECIST criteria for up to two years and thermal-ablation nonresponders (n=24), defined as patients who had progressive disease by localized mRECIST criteria within the two years. Both study cohorts were matched for demographic, clinical baseline and other tumor characteristics as shown in **Table 1**.

### Biopsy collection and FFPE preparation

Needle-biopsy specimen were obtained directly from a Bard Mission 18-gauge spring loaded biopsy gun (BD, Franklin Lakes, NJ)^22^. These 2-cm cores were preserved using a standard protocol^23^.

Specimens were immediately fixed with 10% formalin. Tissues were then dehydrated by 100% ethanol (T038181000, ThermoFisher). Tissues were cleared by 97% xylene (L13317.AP, ThermoFisher). Paraffin was heated to 60°C and allowed to harden overnight. After that, tissue blocks were sectioned in 5μm and mounted on a standard glass slide and covered with a slip cover.

### Proteomic analysis

#### FFPE Sample Extraction & Preparation

290 FFPE slices (n=5 per patient) were digested with trypsin. The supernatants were concentrated by SpeedVac drying and re-dissolving or stored at 4°C until analysis. The sample extract was separated via the mixed mode liquid chromatography with mobile phase gradients. Quality controls were incorporated into each batch to assess and correct for technical effects including QC pooled plasma, and QC water blank samples.

##### Identification

The peak fragmentation from the data dependent MS2 scans were matched to libraries. Moreover, identifications were determined using Thermo Compound Discoverer mzCloud and the internal Dalton software with dot product > 80% and more than 3 matching fragmentation ions.

##### Relative quantification

Acquired raw MS1 data were first aligned using persistent background ions as lock masses. MS1 spectral correlations were then used to align MS1 data retention time to a template run across the entire run period. Background ions were eliminated based on their presence in quality control water samples. Outliers were omitted based on the correlation distance between each sample profile (r < 0.85). Intensity was normalized to correct for variable sample preparation and injection volumes by sample mass.

### Hyperthermic Responsiveness Experiments

HCC SNU423, SNU449, HepG2/C3a cell lines were procured from the American Type Culture Collection (ATCC). Cells were seeded at 1.0x10^5^ cells and cultured in RPMI containing 10% FBS, 4mM L-glutamine, 50Upenicillin/mL, and 50μg streptomycin/mL at 37°C with 5% CO2, for 24 hours. Cell plates were then be placed in a water bath at 37°C or 47°C for 10 minutes and allowed to rest at room temperature for 20 minutes. Cell death was monitored by counting using hemocytometer and 0.4% trypan blue (cat# 15250061, ThermoFisher) at 24, 48, and 72-hour intervals. The cell growth bright-field images were captured using Keyence BZ-X series All-in-One fluorescence microscope.

Additionally, these experiments were performed in SNU449 along with Galectin-1 inhibitor OTX008 (HY-19756, Medchemexpress) and in shGal-1 and respective shControl-SNU449.

### shRNA transfection in SNU449 cells

SNU449 cells were seeded at 3x10^5^ cells in a 12-well plate and cultured in RPMI containing 10% FBS, 4mM L-glutamine, 50Upenicillin/mL, and 50μg streptomycin/mL at 37°C. The following day, cells were transduced with lentiviral shRNA particles carrying three to five expression constructs each encoding target-specific 19-25 nucleotides (plus hairpin) shRNA designed to silence the gene expression of galectin-1 (sc-35441-V, scbt) or respective shControl particles (sc-108080, scbt).

Briefly, 1mL of RPMI mixed with Polybrene (sc-134220, scbt) to get 5 µg/ml, was added into cells. 1x10^6^ lentiviral particles were then slowly added. After 24 hours, the media was replaced with full RPMI containing 10% FBS, 50Upenicillin/mL, and 50μg streptomycin/mL and continued to incubate for another 24 hours. The cells were then split with a 1/3 ratio and continued with incubation for 24 hours. After that, the predetermined dose (15µg/mL) of puromycin (sc-108071, scbt) was used for selection. Successful transfection was validated using western blot. Onwards, cells were selected with 15µg/mL puromycin every 3 days.

### Metabolite Extraction and Metabolomics Analysis of Tumor Lysates

Tumor extraction was performed using our standard metabolite extraction protocol^24^. For data acquisition, tumor metabolite extracts were resuspended in 50% ACN (500μL for the cell extracts, 400μL for the 4 mg tissue extract). The LC separation utilizing an Ion Chromatography System (ICS) 5000 (Thermo Scientific) was performed on a Dionex IonPac AS11-HC-4μm anion exchange column. The gradient was 5-95 mM KOH over 13 min, followed by 5 min at 95 mM KOH, before re-equilibration. Other LC parameters: flow rate 350µl/min, column temperature 35 ⁰C, 10μL injection volume. The Q Exactive mass spectrometer (Thermo Scientific) was operated in negative ion mode for detection of metabolites using a resolution of 70,000 at m/z 200 and a scan range of 70-900 m/z.

Data were analyzed using Maven (version 8.1.27.11). Metabolites were identified based on accurate mass (±5 ppm) and previously established retention times of pure standards. Data analysis was performed using in-house R scripts.

### Western Blotting

Cells were seeded at 4x10^6^ in 10 cm dishes. Cells were then harvested and lysed using the lysis buffer containing 20mM Tris-HCl (15567027, Fisher) with 10μL/mL protease inhibitor cocktail (78440, Fisher). The lysed cells were centrifuged at 17,000g for 5 minutes. Protein concentrations were assessed using Pierce BCA Protein Assay kits (23227, Fisher) along with a SpectraMax M5 microplate reader. The supernatants were heated up to 70°C for 10 minutes. After heating, 30μg protein was used for immunoblotting. Gel was transferred to membrane (LC2003, Fisher) and blocked with Pierce Fast Blocking Buffer (37575, Fisher) for 30 minutes. The primary antibodies with dilution ratio were as follows: galectin-1 1/1000 (Abcam), β-tubulin 1/1000 (15568, Abcam). After primary antibody incubation, membrane was washed with TBST and incubated with secondary antibody goat anti-rabbit IgG conjugated with Horseradish peroxidase (Abcam, ab6721) at 1/5000.

Signals were developed using SuperSignal West Pico (34580, Fisher). Membrane was imaged using 1500 iBright imaging system.

### SNU-449-cell-derived orthotopic implantation

All animal protocols in this study were approved by UCLA. SNU449-cell-derived tumors were orthotopically implanted into the liver following a standard protocol^25^. Briefly, 4-week-old nude mice were anesthetized using isoflurane, and the liver was exposed. A 100mg tumor was attached to the left-lateral lobe of the liver using 5.0 VICRYL suture (NC2872069, Fisher) and wrapped with a 1x0.5-cm piece of SURGICEL (1951, Ethicon). Incisions were closed using wound clips (10-001-024, Fisher). Mice were then administered DMSO or OTX intraperitoneally at 5mg/kg^19^ twice per week for two weeks. Ablation was performed by inserting a 14-gauge antenna connected to ECO Microwave Ablation (MWA) (ECO Inc., China) instrument at 5W for 3 minutes^26^. For sham ablation, the tumors were only inserted with the probe without thermal exposure. After the procedure, mice were continued with DMSO or OTX intraperitoneally at 5mg/kg twice per week for another week. One week following ablation, tumors were harvested for weights, tumor volumes, and metabolomics.

### Statistical Analysis

The log-rank test was utilized to determine the differences in tumor progression-free survival rates^27^. Regarding proteomics, statistical associations between features and experimental variables were tested with multiple linear regression and Benjamin-Hochberg. Unpaired and one-tailed Student’s t-test was used in staining quantification. One-tailed and unpaired Student’s t-test and Chi-square were used for continuous and categorical variables in demographic and clinical characteristics, respectively^28^. P-values less than 0.05 were considered statistically significant. All statistical analysis was performed using Microsoft Excel V16.87.

## V. DISCUSSION

This study investigated the role of Gal-1 in modulating response to thermal ablation via regulating glycolysis. High levels of Gal-1 were identified from a retrospective cohort of pre-ablation biopsy samples of patients who later had localized mRECIST confirmed progressive disease within two years of thermal ablation. Moreover, inhibiting Gal-1, either through gene silencing or pharmacological inhibitors (like OTX008), sensitized the cells to sublethal hyperthermia. Importantly, *in-vivo* studies using an orthotopic murine model showed that the combination of Gal-1 inhibition and thermal ablation led to significantly enhanced tumor reduction compared to thermal ablation alone. Combining Gal-1 inhibition with thermal ablation also led to significantly decreased levels of critical glycolytic metabolites, F-1,6BP, 3PG and PEP along with reduced Gal-1 expression in the tumors. These results underscore Gal-1 as a key player in modulating thermal ablation response via glycolysis.

Metabolomics-based approaches have improved current understanding of cancer biology, especially in solid tumors such as HCC, with known subtypes that are resistance to locoregional and systemic therapies^29,30^. However, efforts to phenotype treatment-resistant HCCs have been limited by historical HCC management guidelines that favor non-invasive imaging for diagnosis rather than tissue sampling^2^. Nevertheless, with the addition of the 2023 AASLD and 2018 EASL guidelines that acknowledge the rapidly expanding role of tissue-based biomarkers, there has been gradual acceptance of the utility of needle biopsies for HCC^2,8^. Increased tissue sampling as part of the patient evaluation will allow for more comprehensive interrogation of the underlying HCC metabolomic tumor microenvironment.

Gal-1 is a glycan-binding protein that has been implicated in the progression of many cancers, with key roles in tumor invasion and treatment resistance^9,10,31,32^. Specifically with HCC, Gal-1 overexpression is particularly notable for its role in regulating the tumor microenvironment and promoting cancer progression^31,33^. Previous research has shown that Gal-1 is overexpressed in HCC tissues compared to normal liver tissues, and its expression has been associated with poor prognosis^31,33^. Gal-1 can influence tumor growth through multiple mechanisms, such as modulating cell adhesion, promoting angiogenesis, and inhibiting T-cell–mediated immune responses^31,33^. However, its role in metabolic regulation, particularly as a response to HCC treatment, has only recently begun to emerge.

Thermal ablation has been well-established to cause immediate cell death at the center of the ablation zone^2,34^. However, the peripheries of the ablation zone are associated with sublethal hyperthermia that can increase rates of aerobic glycolysis in more aggressive subtypes of HCC^17^. Peri-ablation hyperthermia inducing downstream glycolytic products^17,35^ is thus hypothesized to be a key factor contributing to post-treatment resistance in HCC. This study logically builds on prior studies^10,19^ that have also identified Gal-1 as a key player in promoting treatment resistance in HCC. However, while prior studies have focused on the genetic regulatory functions of Gal-1^10,19^, this current study demonstrates metabolic regulation of Gal-1 in the context of peri-ablational hyperthermia. This finding adds a critical dimension to understanding how hyperthermia-induced-metabolic plasticity can contribute to post-ablation HCC recurrence. Gal-1 can thus act as a potential target for therapeutic intervention, particularly in HCC patients that may have a higher risk for progression after thermal ablation.

The *in-vivo* aspect of this study provides key insights into the therapeutic potential of targeting Gal-1 in combination with ablation. This combined strategy led to significantly greater tumor reduction compared to either treatment alone. Moreover, in the combined group, Gal-1 expression was significantly reduced, and glycolysis was disrupted, leading to enhanced tumor cell death. The Gal-1 reduction was likely due to increased activity of the 26S proteasome, which degraded the complex formed by Gal-1 and OTX at a faster speed under hyperthermia^20,21^. Furthermore, the decrease in Gal-1 levels and increased thermal-ablation response in the combined treatment group aligned with our clinical biopsy findings, where Gal-1 expression was lower in responders. These findings underscore the importance of Gal-1 in mediating metabolic adaptation in HCC and suggest that Gal-1 inhibition can effectively enhance the efficacy of thermal ablation.

The findings of this study open several potential avenues for clinical application, particularly in improving patient outcomes. Since Gal-1 plays a central role in facilitating metabolic adaptation to hyperthermic stress, targeting this protein as an adjuvant therapy could make thermal ablation more effective for early-staged HCC. Moreover, the ability of Gal-1 to regulate glycolysis suggests that its inhibition could have broader applications in cancers that exhibit high rates of glycolytic metabolism (i.e., the Warburg effect). Since many solid tumors rely on glycolysis for rapid energy production^13^, Gal-1 inhibitors could be developed as more generalized metabolic disruptors, sensitizing other solid tumors to conventional treatments that induce metabolic stress, such as thermal ablation, radiation, or targeted therapies.

There are several aspects of this study that may limit the generalizability of the results. The biopsy samples from early-stage HCCs may be subject to intra-tumoral heterogeneity. However, we focused on biomarkers that were well conserved across multiple cell types that would mitigate the risks of heterogeneity. Another limitation of the *in-vivo* studies is that the experiments were studied using primarily with a single HCC cell line (SNU449). While an extensive study was performed on several cell types **(Figure S1B)** to isolate the most thermally-resistant HCC subtype, it is unclear if this would extrapolate to other moderately- or poorly-differentiated HCCs. Nevertheless, the thermally-resistant model here was effective in highlighting the potential of Gal-1 inhibition in overcoming ablation-resistance. Future studies will be useful in validating the results in patient-derived HCC cells. Furthermore, extending this to the *in-vivo* study to other mouse models such as hu-BLT mice mouse models would strengthen the generalizability. The focus of Gal-1 on glycolysis, while important for tumor metabolism, may have also inadvertently overlooked other non-glycolytic pathways. For example, Gal-1 has been shown to influence lipid metabolism via hypoxia-inducible factor-1 (HIF-1) dependent pathways in other cancer types^36,37^. Therefore, a more comprehensive metabolomic analysis could provide additional insights into the broader metabolic impact of Gal-1 inhibition.

Post-ablation recurrence for early-stage, non-resectable HCC has been an active area of research, particularly because thermal ablation is applied with curative intent or bridge to transplant^2^. Imaging biomarkers have historically been utilized to identify more aggressive HCC subtypes that may predispose patients to recurrence^38^. However, with the increasing role of HCC biopsy, there are potential tissue biomarkers that can be leveraged for adjuvant therapy. This study provides compelling evidence that Gal-1 plays a critical role in regulating metabolic plasticity in HCC cells, particularly in response to peri-ablational hyperthermia.

Inhibiting Gal-1 can potentially disrupt this metabolic adaptation, sensitizing cancer cells to thermal ablation and enhancing tumor reduction. Characterizing the metabolic role of Gal-1 in post-ablation tumor aggressiveness opens new avenues for therapeutic intervention.

Combining Gal-1 inhibitors with thermal ablation holds great promise for improving patient outcomes, particularly those exhibiting recurrence. While additional research is needed to fully understand the broader metabolic impact of Gal-1 inhibition and to validate these findings in more diverse models, this study represents an important step forward in the development of more effective treatments for early-stage, nonresectable HCC.

