## Supplemental Figures for "Galectin-1 Modulates Hepatocellular Carcinoma Response to Thermal Ablation Through Regulating Glycolysis"

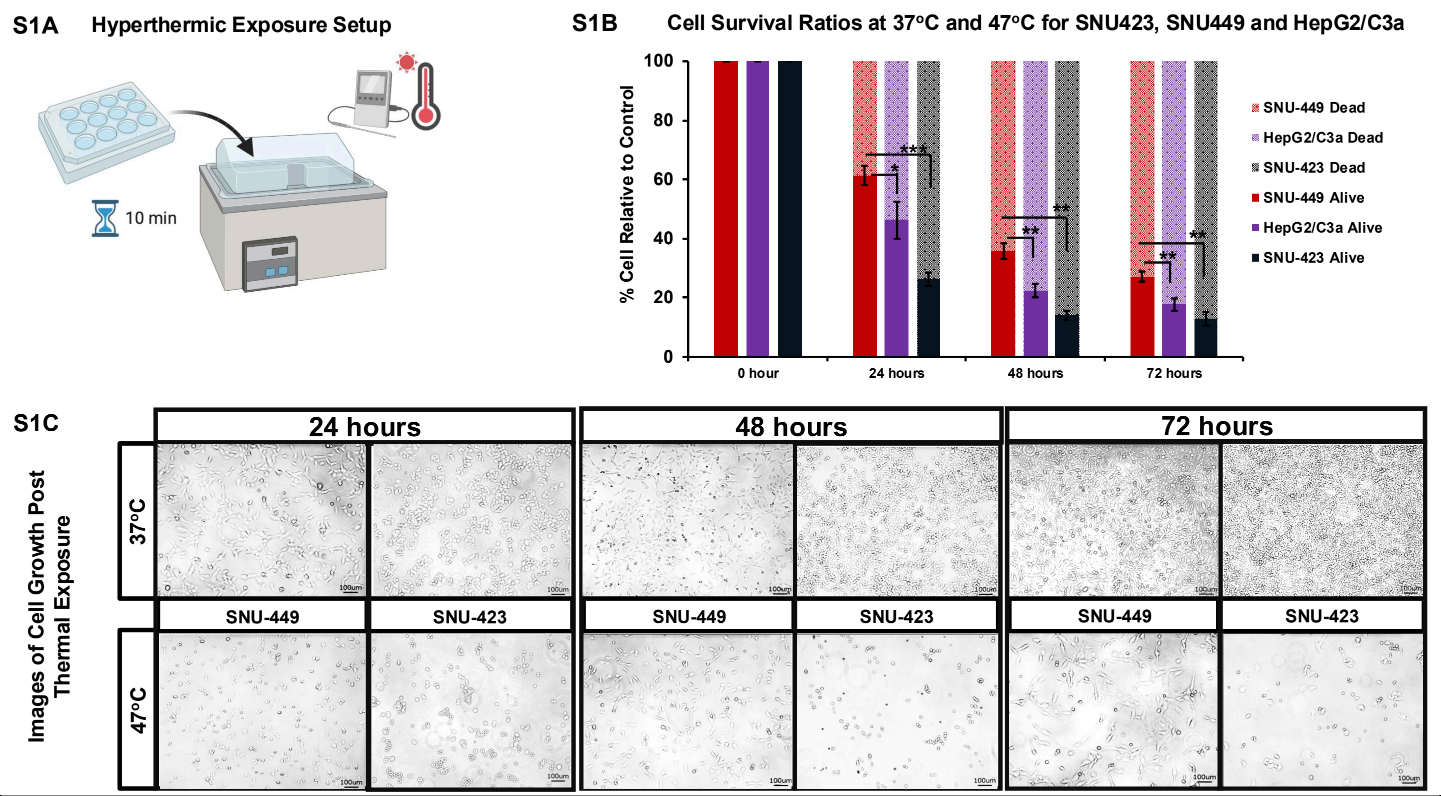


**Figure S1: Hyperthermic responsiveness in HCC cells. A)** Schematic diagram illustrating hyperthermic exposure setup. **B)** SNU449, SNU423, and HepG2/C3a (n=3 each) cell-survival rates post-thermal exposure at 37°C and 47°C at 24, 48, and 72 hours. **C)** Bright-field images of corresponding conditions. p-values were calculated using one-tailed-unpaired student's t-test, *p<0.05, **p<0.01, ***p<0.001.

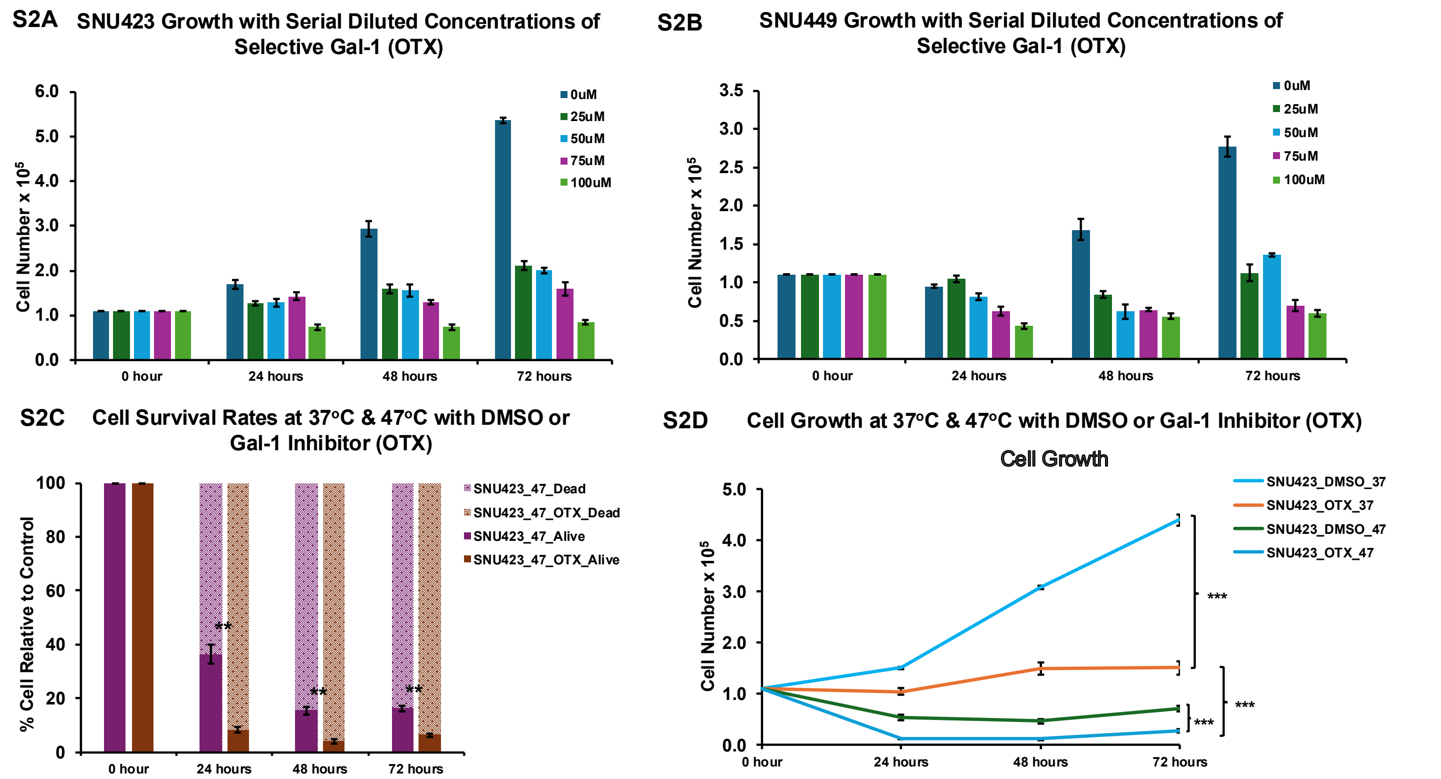


**Figure S2: Decreased galectin-1 levels result in cell death and increased response to hyperthermia. A-B)** SUN449 (A) and SNU423 (B) were treated with galectin-1 inhibitor OTX at various concentrations to examine cell viability. **C)** SNU423 cell-survival rates post thermal exposure at 37°C and 47°C with Gal-1 selective inhibitor OTX or DMSO at 24, 48, and 72 hours. Corresponding cell growth **(D)**. p-values were calculated using one-tailed-unpaired student's t-test, *p<0.05, **p<0.01, ***p<0.001.
